# Engineered α-Synuclein-specific nanobody CAR iTregs restrain neuroinflammation and proteinopathy in Parkinson’s disease mice

**DOI:** 10.64898/2026.08.21.746338

**Authors:** Alice Calderoni, Melania Nannoni, Giorgia Ruffini, Matteo Doglio, Clara Bercher Brayer, Giannelli Sea Serena, Ronald Melki, Monica Casucci, Chiara Bonini, Sharon Muggeo, Vania Broccoli

## Abstract

Parkinson’s disease (PD) is characterized by progressive DAergic neurodegeneration and the accumulation of aggregated α-Synuclein (αSyn), which drives chronic neuroinflammation through sustained activation of innate and adaptive immune responses. Regulatory T cells (Tregs) exert potent immunosuppressive functions and have shown neuroprotective effects in preclinical PD models; however, clinical translation of polyclonal Treg therapies has been limited by poor tissue specificity and insufficient therapeutic efficacy. To overcome these limitations, we engineered induced human Tregs (iTregs) expressing chimeric antigen receptors (CARs) directed against pathological αSyn aggregates. Among the CAR designs tested, only a nanobody-based construct incorporating NbSyn87 displayed selective antigen-dependent activation in response to αSyn preformed fibrils (PFFs). Intriguingly, despite the ability of the parental NbSyn87 nanobody to bind both monomeric and aggregated αSyn, incorporation into the CAR architecture conferred functional selectivity for aggregated conformers. This feature enabled discrimination between pathological extracellular aggregates and physiological monomeric αSyn, providing an important safety advantage. To evaluate therapeutic activity *in vivo*, we established an immunodeficient mouse model of synucleinopathy permissive to human cell engraftment. iTregs preferentially accumulated within αSyn-rich brain regions and, in the presence of astrocyte-derived human IL-2 with antigen-independent mechanism. Conversely, only CAR iTregs directed against αSyn significantly reduced microglial and astrocytic activation, decreased pro-inflammatory cytokine expression, and attenuated αSyn pathology. Collectively, these findings demonstrate that αSyn-specific CAR iTregs can selectively exert potent local immunomodulatory effects, establishing a promising antigen-specific cellular immunotherapy platform for PD and other synucleinopathies.

## Introduction

Parkinson’s Disease (PD) is the second most common neurodegenerative disorder, clinically characterized by motor impairment and pathologically defined by the loss of DAergic neurons in the *substantia nigra pars compacta* (SNpc) (Kalia and Lang, 2015; Bloem et al., 2021). Central to this process is the intraneuronal accumulation of α-Synuclein (αSyn) insoluble oligomers and fibrils which interfere with axonal transport, mitochondrial integrity, and synaptic function, ultimately triggering a cascade of selective neuronal cell loss (Lee and Trojanowski, 2006; Poewe et al., 2017). Upon neuronal death, the release of misfolded αSyn species into the extracellular space acts as a potent Damage-Associated Molecular Pattern (DAMP). These extracellular fibrils engage with Pattern Recognition Receptors (PRRs), such as Toll-like receptor 2 (TLR2), on the surface of resident microglia, initiating a robust innate immune response (Kim et al., 2013). This chronic activation results in the sustained secretion of pro-inflammatory mediators, including TNFα, IL-1β, and reactive oxygen species (ROS), which further exacerbates the neurodegenerative environment (Bido et al., 2021; Geng et al., 2025). This self-perpetuating cycle of protein aggregation and neuroinflammation suggests that the inflammatory reaction is no longer considered a sterile consequence of neuronal death but a co-primary driver of pathology. Peripheral immune profiling of PD patients reveals expansion of pro-inflammatory T cell subsets and impaired immune regulation. CD4⁺ T cells shift toward Th1 and Th17 phenotypes, producing IFNγ and IL-17, which enhance microglial activation and neuronal injury (Tansey et al., 2022; Roodveldt et al., 2024). CD8^+^ T cells infiltrate the SNpc early in PD and may directly mediate DAergic neuron cytotoxicity (Brochard et al, 2009; Galiano-Landeira et al., 2020). These adaptive immune changes correlate with disease progression, suggesting that peripheral immune dysregulation contributes to the central nervous system (CNS) pathology (Terkelsen et al., 2022). Furthermore, genome-wide association studies (GWAS) have linked HLA-DR polymorphisms to PD risk, reinforcing the hypothesis that an aberrant adaptive immune response to αSyn is central to the disease (Nalls et al., 2014). Regulatory T cells (Tregs) are a specialized subpopulation of CD4^+^ T lymphocytes, characterized by the expression of the transcription factor FOXP3, which serve as indispensable gatekeepers of immune homeostasis and peripheral self-tolerance (Sakaguchi et al., 2008). Their primary physiological role involves the suppression of aberrant immune responses and the limitation of chronic inflammation, primarily through the secretion of anti-inflammatory cytokines such as IL-10 and TGFβ, and via direct cell-to-cell contact mechanisms (Josefowicz et al., 2012). In neuroinflammatory contexts, Tregs have emerged as critical mediators of neuroprotection due to their ability to modulate the local innate immune environment (Schwartz and Deczkowska, 2016). However, in PD patients, the endogenous Treg pool often appears numerically diminished or functionally exhausted, failing to counteract the inflammatory milieu (Saunders et al., 2012; Kustrimovic et al., 2018). This decline in regulatory potency prevents the effective dampening of the neuroinflammatory environment within the SNpc, thereby permitting the uncontrolled progression of DAergic neuronal loss. For instance, studies have demonstrated that Tregs isolated from PD patients show a reduced capacity to suppress effector T cell proliferation compared to healthy age-matched controls, suggesting that the systemic immune dysregulation in PD is not merely a byproduct of neurodegeneration but a contributing driver of the disease (Saunders et al., 2012). While the global enhancement of regulatory T cell activity aims to quench neuroinflammation, it simultaneously risks compromising the host’s physiological immune surveillance, potentially increasing vulnerability to opportunistic infections and diminishing anti-tumor responses (Sakaguchi et al., 2008). Recent investigations have highlighted that inside the CNS exogenously administered conventional Tregs exhibit reduced persistence. Without sustained, antigen-specific activation, traditionally mediated by TCR stimulation, and adequate local trophic support (such as IL-2), these cells suffer from lineage instability, downregulating FOXP3 expression (Vahl et al. 2014, Fan et al. 2018). Consequently, they succumb to the hostile inflammatory and oxidative microenvironment, characterized by excessive production of reactive oxygen and nitrogen species, pro-inflammatory cytokines, and metabolic stress, highlighting the need for targeted receptor engineering to ensure long-term functional persistence.

CAR-mediated antigen engineering can empower Tregs to deliver site-specific immunomodulation, thereby ensuring a more targeted approach to neuroprotection (Freitag et al., 2020; Raffin et al., 2020). Upon antigen binding, the CAR structure, comprised of a flexible hinge, a transmembrane domain, and an intracellular signaling tail integrating stimulatory and co-stimulatory domains, transmits a robust activation signal that preserves FOXP3 stability and enhances suppressive fitness (Dawson et al., 2020; Rosado-Sánchez et al., 2023). In this study, we developed a novel immunotherapeutic strategy by engineering human CD4^+^ T cells to express both the transcription factor FOXP3 and a second-generation CAR specific for αSyn. This approach redirects the suppressive power of Tregs specifically toward αSyn aggregates to curb neuroinflammation. In fact, by tethering Treg effector functions directly to the site of neurodegeneration, these engineered cells facilitate localized bystander suppression, effectively quenching the inflammatory cascade within the SNpc while mitigating the deleterious systemic consequences and lineage instability inherent to conventional polyclonal therapies.

## Results

### Generation of stable and functional αSyn-CAR iTregs

We generated a lentiviral (LV) vector encoding for human *FOXP3* gene together with a second-generation CAR construct, constituted by an antigen binding domain fused to a spacer derived from the extracellular domain of the human low-affinity nerve growth factor receptor (NGFR), a CD28 transmembrane domain, the CD3ζ stimulatory domain, and the intracellular signaling domain of CD28, which has been shown to be optimal for Treg phenotype and function (Boroughs et al., 2019). (LV:αSyn-CAR.28z) (Figure 1A). For αSyn recognition, we considered two different antigen binding domains: a single-chain variable fragment (scFv) and a nanobody. The αSyn scFv was engineered from the variable heavy (VH) and variable light (VL) domains of the 306C7B3 antibody, which binds aggregated αSyn species with high selectivity and picomolar affinity (Duchs et al., 2023). The VH and VL domains were connected by the Whitlow linker (Whitlow et al., 1993). Alternatively, the NbSyn87 nanobody was chosen for its ability to recognize a C-terminal epitope of αSyn with high affinity, binding both monomeric and aggregated αSyn species (Guilliams et al., 2013) (Figure 1A). Finally, an anti-CD19 CAR construct, identical in design to the above-described vectors, but targeting an unrelated antigen, was used as a negative control (Doglio et al., 2024) (Figure 1A). To generate αSyn-CAR iTregs, CD4^+^ T cells were isolated from peripheral blood of healthy donors and stimulated in the presence of αCD3/28 coated beads (Figure 1B). After LV transduction, cells were expanded in the presence of high IL-2 and rapamycin and showed robust expression of the NGFR marker embedded into the CAR constructs and FOXP3, as confirmed by flow cytometry and RT-qPCR, while only Nb-αSyn-CAR expressed NbSyn87 as expected (Figures 1C,D). Mean transduction efficiency, assessed by NGFR and FOXP3 expression by flow cytometry, was 76.9% ± 5.9%, 62.5% ± 8.4%, 76.5% ± 5.8% for LV:Nb-αSyn-CAR, LV:scFv-αSyn-CAR and LV:CD19-CAR respectively, with stable co-expression of the transgenes maintained until the last time point analyzed (21 days in culture) (Figure 1C). Droplet digital PCR (ddPCR) analysis showed a mean vector copy number (VCN) of 5.76 ± 1.18, 2.38 ± 0.13, 3.73 ± 0.94 per cell, in LV:Nb-αSyn-CAR, LV:scFv-αSyn-CAR and LV:CD19-CAR transduced cells, respectively (Figure 1E). Nb-αSyn-CAR iTregs, scFv-αSyn-CAR iTregs and CD19-CAR iTregs displayed a similar expansion rate (29.3% ± 4.2%, 34.4% ± 4.2% and 33.6% ± 5.64% for NB, scFv and CD19 respectively) at day 9 (Figure 1F), indicating that the lentiviral transduction did not affect CD4^+^ T cell expansion. All the αSyn-CAR iTregs displayed a Treg-like phenotype respect to untransduced cells, characterized by the upregulation of immune checkpoints and co-signaling markers including CD25, FOXP3, GARP, HELIOS and CTLA4, assessed by flow cytometry (Figure 1G) and TIGIT, GITR, ICOS, HLA-DRA, and IL2RA (CD25) as confirmed by RT-qPCR (Figure 1H). Collectively, these findings demonstrate that CAR iTregs can be readily generated and expanded, while maintaining an unaltered proliferation kinetics with stable phenotype.

**Figure 1.**
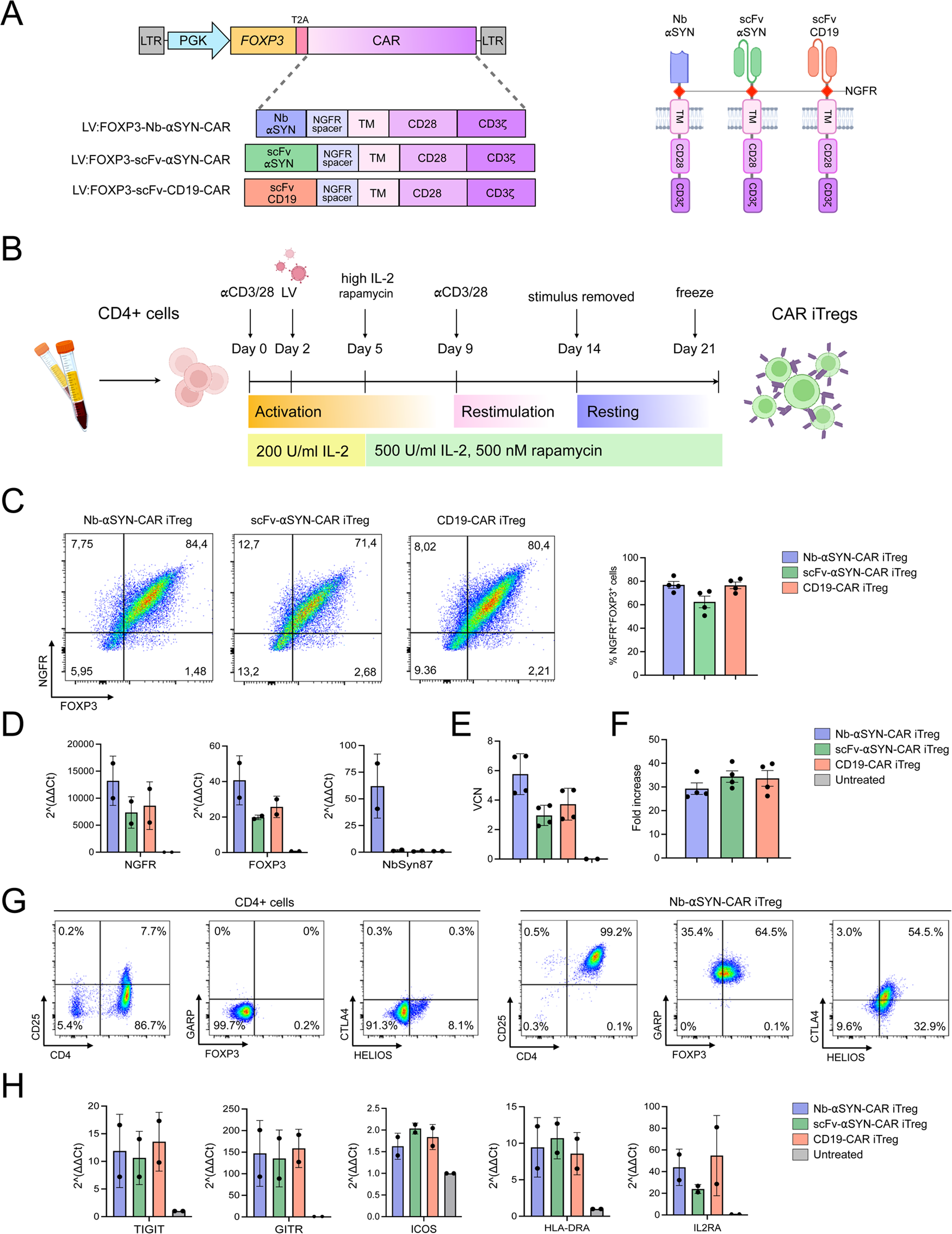
Generation and characterization of CAR iTreg. **A)** Left: Schematic representations of LVs containing the FOXP3 gene and different chimeric antigen receptors (CARs). Right: illustration of the CAR protein domain architectures. **B)** Schematic representation of the experimental timeline used for activation and transduction of primary CD4⁺ T cells and expansion of CAR iTregs, prior to functional assays. **C)** Representative flow cytometry plots and corresponding quantification of efficiency of transduction, n=4. **D)** Transgenes expression levels in engineered CAR iTregs compared to untransduced cells, n=2. **E)** Vector copy number (VCN) analysis per genome in transduced CAR iTregs, n=4. **F)** Fold increase in CAR iTregs expansion at day 9 of culture, n=4. **G)** Representative flow cytometry of the typical Treg markers s CD25, GARP, FOXP3, CTLA4 and HELIOS in untreated CD4^+^ T cells and Nb-αSyn-CAR iTregs. **H)** Gene expression analysis (2^−ΔΔCt^ relative to untransduced cells) of key Treg-related markers and cytokines across the different CAR iTreg constructs, n=2. Data are presented as mean ± SEM. Statistical significance was determined using ∗ *P* < 0.05, ∗∗ *P* < 0.01, ∗∗∗ *P* < 0.001, ∗∗∗∗ *P* < 0.0001, ns *P* > 0.05.

### Antigen-specific selectivity of the αSyn binders in brain tissue

Given the feasibility of generating CAR constructs incorporating αSyn-binding domains, it was essential to validate their binding specificity and assess the background reactivity of the αSyn-specific scFv and nanobody used in the CAR on tissue sections. For this goal, we fused the human IgG Fc domain to both binders to enable their recognition with a secondary antibody coupled to a fluorescent reporter for immunofluorescence analysis. Then, sections from mouse brains previously injected with LV:SNCA-A53T (hereafter LV:SNCA) and harvested 8 weeks post-transduction, were stained with these Fc-fused binders. To assess the ability of these binders to recognize different αSyn protein species, sections were simultaneously co-stained with the antibody against pS129αSyn, a hallmark of αSyn aggregated species, and a second antibody recognizing both human αSyn protein monomers and aggregates (Bido et al., 2021) (Figure 2). Both scFv- and Nb-derived binders exhibited highly selective immunoreactivity within the human αSyn overexpressing areas, with no detectable staining in healthy mouse brain tissue (Figure 2). Moreover, high-magnification imaging revealed that the αSyn nanobody signal was enriched in regions containing the monomeric αSyn protein (Figure 2, arrows). In contrast, the αSyn scFv signal was predominantly detected in areas exhibiting strong pS129αSyn immunoreactivity, indicating selective localization to pathological αSyn aggregates (Figure 2, arrowheads). Collectively, these findings indicate that both binding domains efficiently recognize human αSyn *in vivo*, supporting their use as the extracellular recognition modules of our CAR Treg constructs. Notably, whereas the αSyn scFv preferentially recognizes pathological αSyn aggregates, the αSyn nanobody lacks protein aggregate specificity and binds both monomeric and aggregated αSyn.

**Figure 2.**
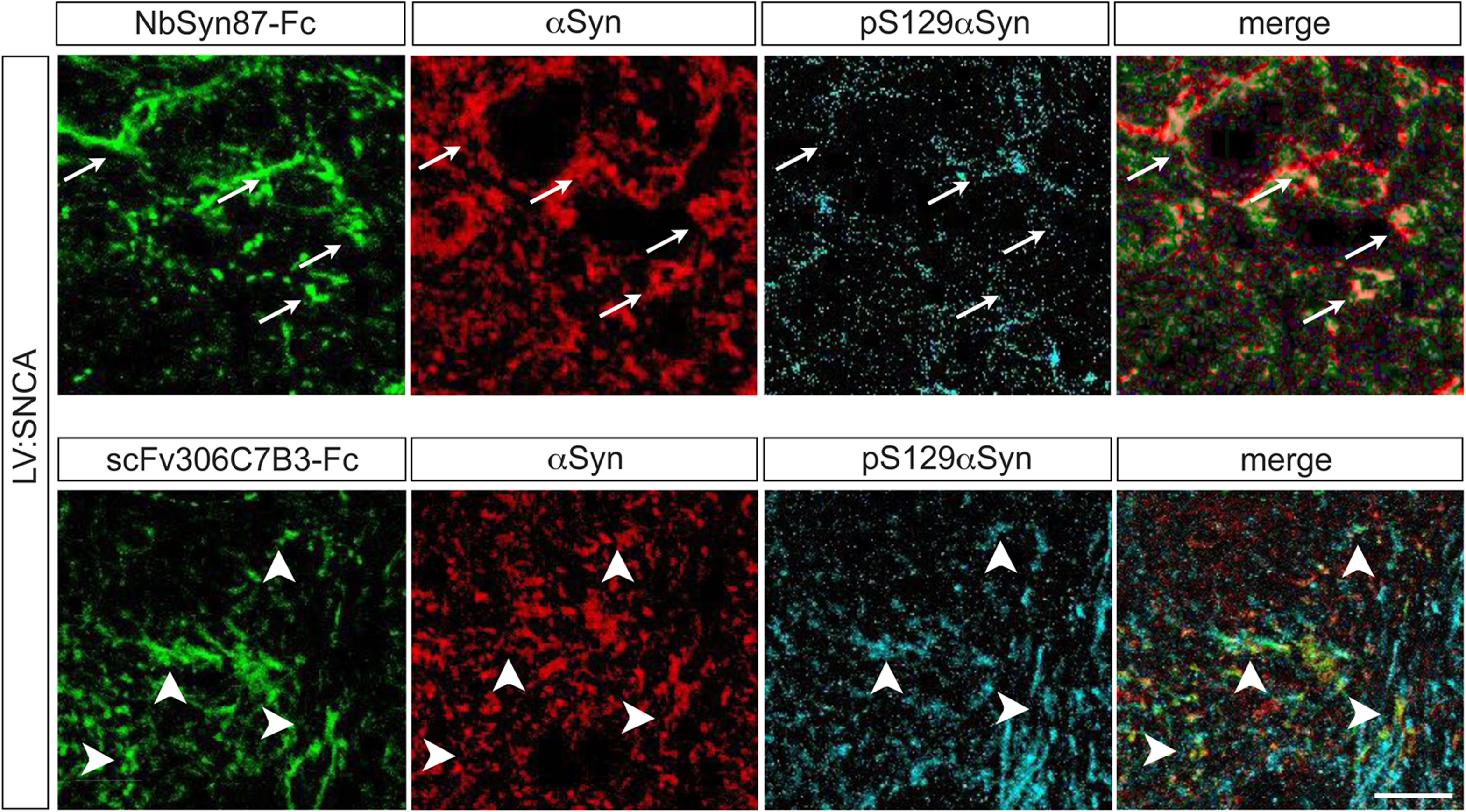
NbSyn87 and scFv306C7B3 exhibit distinct *in vivo* binding selectivity toward αSyn protein species. Representative immunofluorescence images showing NbSyn87 or scFv306C7B3 (green) together with total αSyn (red) and phosphorylated αSyn (pS129-αSyn; cyan) in the *substantia nigra*. Arrows indicate the preferential localization of NbSyn87 to regions enriched in monomeric αSyn, whereas arrowheads highlight the predominant localization of scFv306C7B3 within pS129αSyn-positive aggregates. Images are representative of three independent biological replicates. Scale bar: 30 µm.

### Antigen-specific activation of anti αSyn-CAR iTregs by αSyn fibrils

We next evaluated whether engineered CAR iTregs could be activated using an antigen-specific stimulus by an *in vitro* proliferation assay (Figure 3A). For this goal, we employed the αSyn pre-formed fibrils (PFFs), that have been extensively demonstrated, including in our previous studies, to efficiently seed synucleinopathy pathology in both cells and mouse brains (Iannielli et al., Rey et al, 2019; Srivastava et al., 2020). Nb-αSyn-CAR iTregs exhibited robust and selective activation-induced proliferation upon exposure to αSyn PFFs, while maintaining a quiescent state in the absence of the target antigen (Figure 3B). Conversely, αSyn monomers failed to induce activation, highlighting the pronounced selectivity of these cells for aggregated αSyn species such as PFFs. In contrast, CD19-CAR iTregs activation was achieved only with the polyclonal stimulus exerted by αCD3/28, but not in the presence of αSyn (Figure 3B). Furthermore, Nb-αSyn-CAR iTregs exhibited dose-dependent activation in response to decreasing concentrations of αSyn PFFs, with detectable reactivity observed at femtomolar concentrations (Figure 3C).

**Figure 3.**
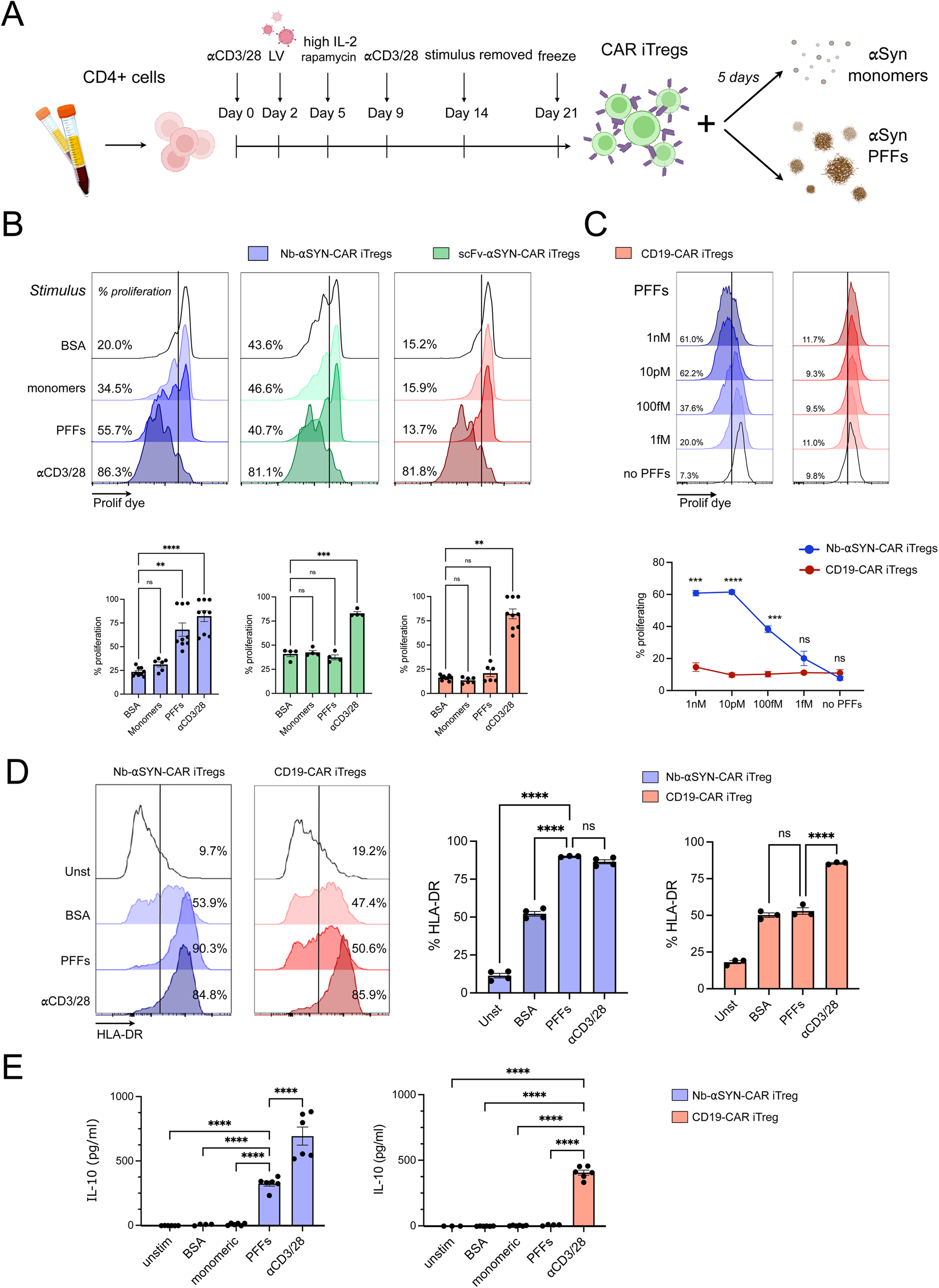
αSyn fibrils promote antigen-specific activation of anti αSyn-CAR iTregs. **A)** Schematic representation of the experimental workflow used for *in vitro* activation assays. **B)** Representative proliferation assessment of CAR iTregs by flow cytometry upon different stimuli (top), with corresponding quantifications (bottom), n=4-9. Data are presented as mean ± SEM. Statistical significance was calculated using one-way ANOVA followed by Tukey’s multiple comparisons test. ∗∗ *P* < 0.01, ∗∗∗ *P* < 0.001, ∗∗∗∗ *P* < 0.0001., ns *P* > 0.05. **C)** Dose-dependent proliferation response of CAR iTregs exposed to decreasing concentrations of PFFs. Representative histograms (top), and dose-response curves (bottom), n=3. Data are presented as mean ± SEM. Statistical significance was calculated using one-way ANOVA followed by Tukey’s multiple comparisons test. ∗∗∗ *P* < 0.001, ∗∗∗∗ *P* < 0.0001, ns *P* > 0.05. **D)** Cell surface expression of the activation marker HLA-DR on CAR iTregs following antigen stimulation. Representative flow cytometry histograms (left), and corresponding bar graphs (right), n=3-4. Data are presented as mean ± SEM. Statistical significance was calculated using one-way ANOVA followed by Tukey’s multiple comparisons test. ∗∗∗∗ *P* < 0.0001, ns *P* > 0.05. **E)** Quantification of IL-10 secretion (pg/ml) in culture supernatants of CAR iTregs activation assays, n=3. Data are presented as mean ± SEM. Statistical significance was calculated using one-way ANOVA followed by Tukey’s multiple comparisons test. ∗∗∗∗ *P* < 0.0001.

Unexpectedly, flow cytometry analysis revealed that scFv-αSyn-CAR iTregs exhibited similarly moderate rates of cell proliferation in all tested conditions regardless of the presence of αSyn (Figure 3B). This phenomenon, characteristic of tonic signaling, suggests that scFv-mediated surface clustering induces an undesired, persistent state of constitutive activation that was shown to ultimately compromise Treg stability (Barden et al, 2024). To investigate whether the constitutive activation observed in scFv-αSyn-CAR iTregs was driven by spontaneous receptor aggregation, we assessed their surface distribution. Leveraging the NGFR-derived spacer for direct detection, confocal imaging demonstrated that the scFv construct exhibited distinct receptor clustering on the cell membrane, unlike the homogeneous distribution observed in the other constructs (Figure S1). Notably, this spatial aggregation strongly correlates with the non-specific activation observed in our functional assays, validating that the baseline tonic signaling stems from spontaneous physical clustering. These data demonstrate that the Nb-based CAR successfully confers antigen-specific signaling, allowing the cells to sense the pathological protein aggregates characteristic of PD, while avoiding the detrimental effects of constitutive, ligand-independent signaling. To confirm functional responsiveness, we performed an activation assay evaluating HLA-DR upregulation and IL-10 secretion on CAR iTregs. Notably, Nb-αSyn-CAR iTregs exhibited specific activation upon exposure to PFFs (Figure 3D). Conversely, CD19-CAR iTregs were activated at high levels only by polyclonal stimulation, while their response to PFFs remained comparable to that of unrelated antigen (Figure 3D). Consistently, Nb-αSyn-CAR iTregs released high levels of IL-10 selectively upon PFF engagement (Figure 3E). In contrast, IL-10 secretion by CD19-CAR iTregs was strictly restricted to polyclonal activation, with PFFs exposure eliciting no detectable response (Figure 3E). Together, these data confirm that CAR iTreg activation and cytokine release are strictly antigen specific.

### Antigen-specific suppressive functions of Nb-αSyn-CAR iTregs

We then proceeded to assess the suppressive functions of CAR iTregs after exposure to polyclonal or antigen-specific stimulation and their effects on conventional CD4^+^ T cells (Figure 4). After 3 days of co-culture between iTregs and CD4^+^ T cells exposed to a polyclonal stimulation with αCD3/CD28 beads, we measured proliferation rates of the latter cell type by flow cytometry (Figure 4A). Both Nb-αSyn CAR iTregs and CD19-CAR iTregs effectively inhibited conventional CD4^+^ T cells proliferation in a dose-dependent manner. (Figure 4B). We observed a comparable suppressive capacity between Nb-αSyn-CAR and CD19 CAR iTregs, with a suppression index of 72.3% ± 4.4% and 68.8% ± 4.0% at the highest ratio, respectively, while conventional T cells exerted no suppression abilities, as expected (Figure 4B). This suppressive effect was further validated by a significant increase of the immunomodulatory cytokine IL-10 (maximal concentration 7.8 ± 0.8 and 6.0 ± 0.5 ng/ml for Nb-αSyn-CAR iTregs and CD19-CAR iTregs, respectively) with an overlapping dose-response kinetics between Nb-αSyn-CAR iTregs and CD19-CAR iTregs consistent with their induced regulatory function (Figure 4C). These findings indicate that both αSyn- and CD19-CAR iTregs shared a similar response to a polyclonal stimulus developing comparable immunosuppressive functions. Next, we assessed whether CAR iTregs were responsive to an antigen-specific stimulation. For this goal, CAR iTregs were primed with αSyn PFFs for two days, while conventional CD4^+^ T cells were activated with αCD3/CD28 beads for 24 h. Both cell types were then co-cultured in the presence of αSyn PFFs (Figure 4D). In presence of PFFs stimuli, Nb-αSyn-CAR iTregs, but not anti CD19-CAR iTregs, suppressed cell proliferation (34.9% ± 2.9% vs 7.8% ± 3.7% of suppression at the highest ratio, respectively) (Figure 4E), highlighting the suppressive efficacy of the αSyn-CAR in an antigen-specific manner. Concordantly, only Nb-αSyn-CAR iTregs, but not CD19-CAR iTregs, showed enhanced IL-10 levels in a dose dependent manner in the co-colture supernatants in presence of αSyn PFFs (Figure 4F). Altogether, these findings confirm that our Nb-αSyn-CAR engineering approach not only confers specific response of iTregs toward αSyn aggregated species but also enhances their suppressive functions specifically in the presence of the target antigen.

**Figure 4.**
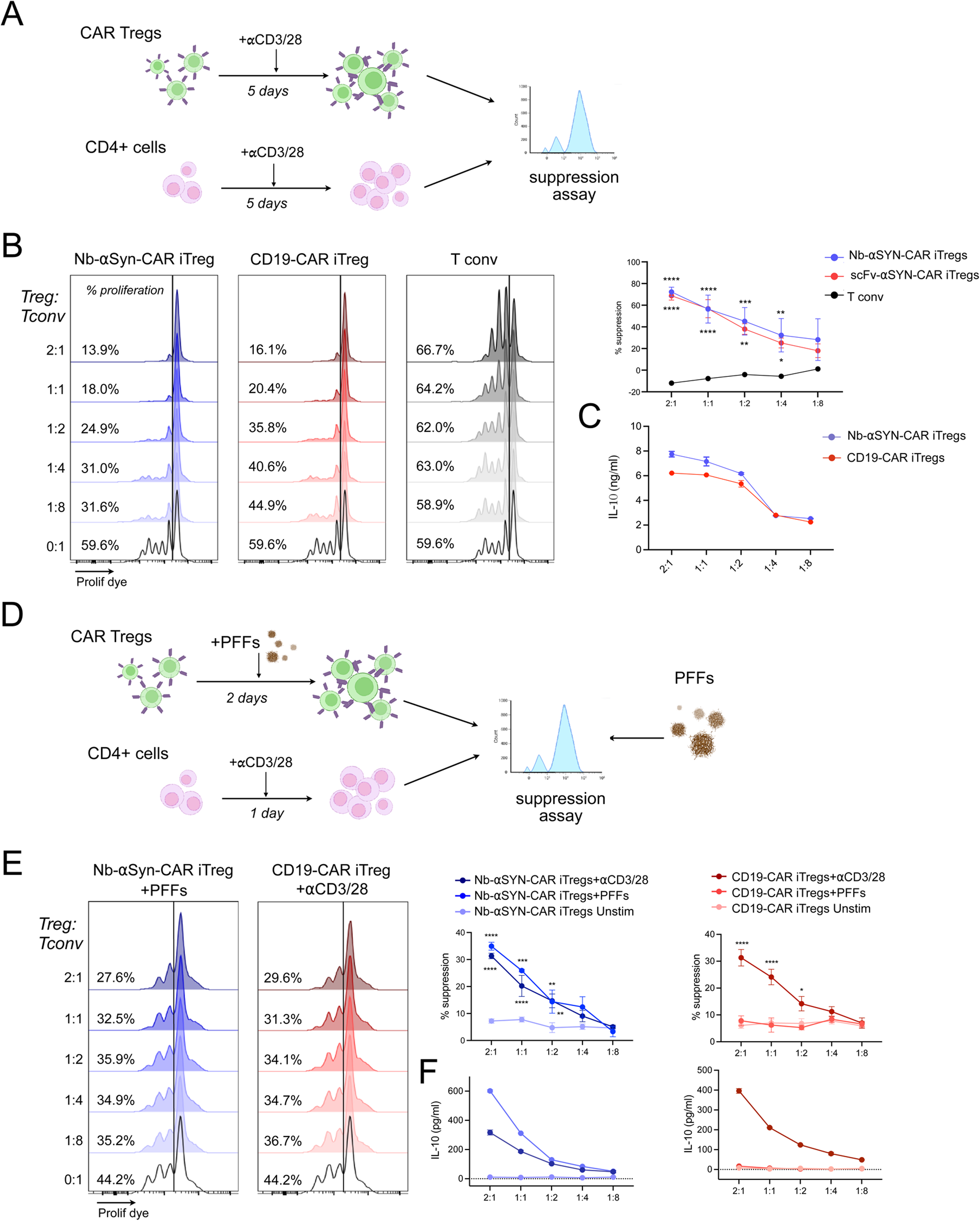
Nb-αSyn- and CD19-CAR iTregs exhibit shared polyclonal suppression but distinct antigen-driven responses. **A)** Schematic representation of the experimental workflow used for *in vitro* polyclonal suppression assays. **B)** Polyclonal suppression of Tconv proliferation across decreasing Treg:Tconv ratios (from 2:1 to 1:8). Representative flow cytometry histograms (left), and corresponding percentage of suppression curves (right), n=3. Data are presented as mean ± SEM. Statistical significance was calculated using one-way ANOVA followed by Tukey’s multiple comparisons test. ∗ *P* < 0.05, ∗∗ *P* < 0.01, ∗∗∗ *P* < 0.001, ∗∗∗∗ *P* < 0.0001. **C)** Quantification of IL-10 secretion (ng/ml) in culture supernatants of CAR iTregs polyclonal suppression assays, n=3. **D)** Schematic representation of the antigen-specific suppression assay workflow. **E)** Antigen-dependent suppressive capacity of CAR iTregs across decreasing Treg:Tconv ratios (from 2:1 to 1:8). Representative histograms (left), and corresponding percentage of suppression curves (right), n=3. Data are presented as mean ± SEM. Statistical significance was calculated using one-way ANOVA followed by Tukey’s multiple comparisons test. ns = not significant, ∗ *P* < 0.05, ∗∗ *P* < 0.01, ∗∗∗ *P* < 0.001, ∗∗∗∗ *P* < 0.0001. **F)** Quantification of IL-10 secretion (pg/ml) in culture supernatants of CAR iTregs antigen specific suppression assays, n=3. Data are presented as mean ± SEM. Statistical significance was calculated using one-way ANOVA followed by Tukey’s multiple comparisons test. ∗ *P* < 0.05, ∗∗ *P* < 0.01, ∗∗∗ *P* < 0.001, ∗∗∗∗ *P* < 0.0001., ns *P* > 0.05.

### Nb-αSyn-CAR iTregs suppress the activation state of human myeloid-induced microglia-like cells

We then sought to investigate the immunomodulatory activity of Nb-αSyn-CAR iTregs. To this end, we leveraged an established differentiation platform that generates human induced microglia-like cells (iMGCs) from peripheral blood myeloid cells after 14 days of culture with GM-CSF and IL-34 (Figure 5A) (Ohgidani et al., 2014). These cells recapitulate features of human microglia, including a plastic morphology, expression of canonical microglial markers, phagocytic competence, and rapid responsiveness to inflammatory stimuli, making them a valuable *in vitro* model for studying neuroinflammatory mechanisms and therapeutic immunomodulation (You et al., 2023). Following cytokine exposure, cells were characterized by high TMEM119, CD14, HLA-DR, CD86, CD206, and CD163 expression. This population co-expressed markers typically associated with both classically activated (M1-like; HLA-DR and CD86) and alternatively activated (M2-like; CD163 and CD206) microglia/macrophages, indicating the acquisition of a mixed activation state under these culture conditions (Ransohoff, 2016) (Figure 5B). These cells displayed a variable and dynamic morphology in culture while uniformly expressing the myeloid marker IBA1 (Figure 5C). Next, Nb-αSyn-CAR iTregs previously activated with PFFs or the αCD23/28 polyclonal stimulus were co-cultured with iMGCs for 24 hrs before exposing them to the pro-inflammatory cytokines IFNγ and LPS (Figure 5A). In parallel, iMGCs were cultured in the conditioned medium of Nb-αSyn-CAR iTregs activated with PFFs (Figure 5A). After 6 hrs from the pro-inflammatory cues iMGCs were profiled by qRT-PCR and after 24 hrs by flow cytometry. Intriguingly, cytokine-induced upregulation of the pro-inflammatory marker CD86 was markedly attenuated in the presence of Nb-αSyn-CAR iTregs or their conditioned medium (Figure 5D). Conversely, the proportion of cells expressing the pro-resolving marker CD206 was significantly increased under the same conditions (Figure 5D). Consistent with this phenotypic shift, Nb-αSyn-CAR iTregs broadly suppressed cytokine-induced activation of iMGCs, as evidenced by the marked downregulation of multiple inflammatory mediators, including IL1B, TNFA, IL6, NLRP3, and CCL2 (Figure 5E). Notably, SOCS3 expression was significantly lower in the presence of Nb-αSyn-CAR iTregs than in cells treated with their conditioned medium alone (Figure 5E). Given that SOCS3 is predominantly induced by soluble mediators, such as IL-10, via the JAK/STAT3 signaling pathway (Kubo et al., 2003), the stronger induction observed with the conditioned medium may reflect its higher concentration of secreted immunomodulatory factors compared with the direct co-culture conditions. Collectively, these findings indicate that Nb-αSyn-CAR iTregs effectively suppress the inflammatory activation of myeloid-derived microglia-like cells, resulting in broad inhibition of multiple pro-inflammatory signaling pathways.

**Figure 5.**
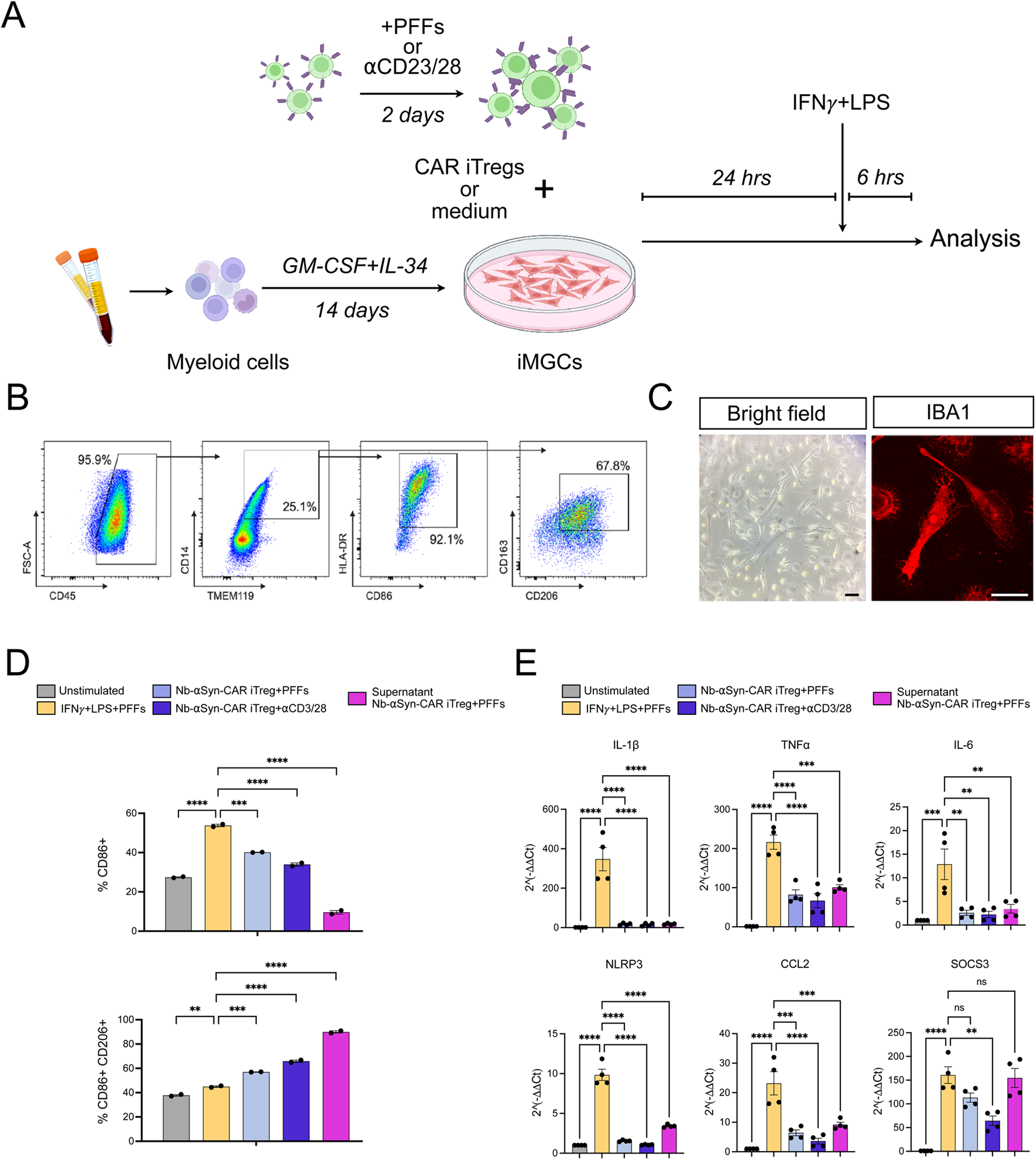
Nb-αSyn-CAR iTregs suppress the inflammatory response of human myeloid-induced microglia-like cells (iMGCs). **A)** Schematic representation of the co-culture setup. Induced microglial-like cells (iMGCs) were differentiated from peripheral blood myeloid cells using GM-CSF and IL-34. iMGCs were co-cultured with Nb-aSyn-CAR iTregs (pre-stimulated with PFFs or aCD3/28) or treated with CAR iTreg supernatant for 24 hours prior to stimulation with IFNg + LPS for 6 or 24 hours before analysis. **B)** Representative gating strategy used to characterize iMGCs based on CD45, TMEM119, CD14, HLA-DR, CD86, CD163, and CD206 expression. **C)** Representative bright-field image (left) showing morphology of iMGCs and immunofluorescence staining (right) for the microglial marker IBA1. Scale bars: 50 µm. **D)** Flow cytometry analysis showing the percentage of % CD86^+^ (top) and % CD86^+^CD206^+^ (bottom) iMGCs across different experimental conditions. n=2. Data are presented as mean ± SEM, statistical significance was determined using one-way ANOVA followed by Tukey’s multiple comparisons test. **P < 0.01, ***P < 0.001, ****P< 0.0001. **E)** Quantitative RT-PCR analysis of pro-inflammatory and suppressive gene expression levels normalized to 18S. n=4. Data are presented as mean ± SEM, statistical significance was determined using one-way ANOVA followed by Tukey’s multiple comparisons test. **P < 0.01, ***P < 0.001, ****P < 0.0001.

### Local hIL-2 expression sustains CAR iTregs homing and survival in the brain

To establish a robust humanized preclinical model of PD, NSG (NOD scid gamma) mice were utilized to prevent xenogeneic rejection of human donor cells. This immunodeficient strain was stereotaxically injected in SNpc with a LV constitutively and ubiquitously overexpressing the human αSyn encoding gene SNCA bearing the A53T mutation (Figure 6A). The A53T missense mutation is highly clinically relevant, as it is linked to early-onset, familial PD and significantly accelerates the kinetics of αSyn aggregation and fibrillization compared to the wild-type protein (Polymeropoulos et al., 1997; Giasson et al., 2002). Treated mice exhibit robust histopathological features of PD characterized by marked αSyn accumulation within the nigral tissue accompanied by a pronounced neuroinflammatory response (Bido et al., 2024). To investigate the homing and engraftment capacity of engineered CAR iTregs within the diseased brain microenvironment, Nb-αSyn-CAR iTregs cells were intravenously administered via the tail vein 8 weeks after the initial LV delivery. However, analysis failed to detect CAR iTregs within the brain parenchyma at 1 week post-infusion, suggesting that the lack of supportive niche or trophic factors severely compromised their *in vivo* survival. To overcome this limitation, we co-injected an adeno-associated virus (AAV) (1×10^7^ vg/site) encoding human interleukin-2 (IL-2) and NGFR under an astrocyte-specific promoter (AAV-GFAP-IL2-NGFR) alongside the LV administration. The localized expression of the transgene was confirmed by immunofluorescence revealing a robust and specific NGFR signal colocalizing with GFAP-positive cells (Figure S2A). Nb-αSyn-CAR iTregs were then administered either as a single infusion or as two infusions given 6 days apart, and their persistence in the nigral tissue was evaluated 9 days after the final dose. Whereas only a small number of CAR iTregs were detected following a single infusion, repeated administration increased their abundance approximately threefold (Figure S2B). Based on these findings, the repeated dosing regimen was selected for all subsequent experiments.

**Figure 6.**
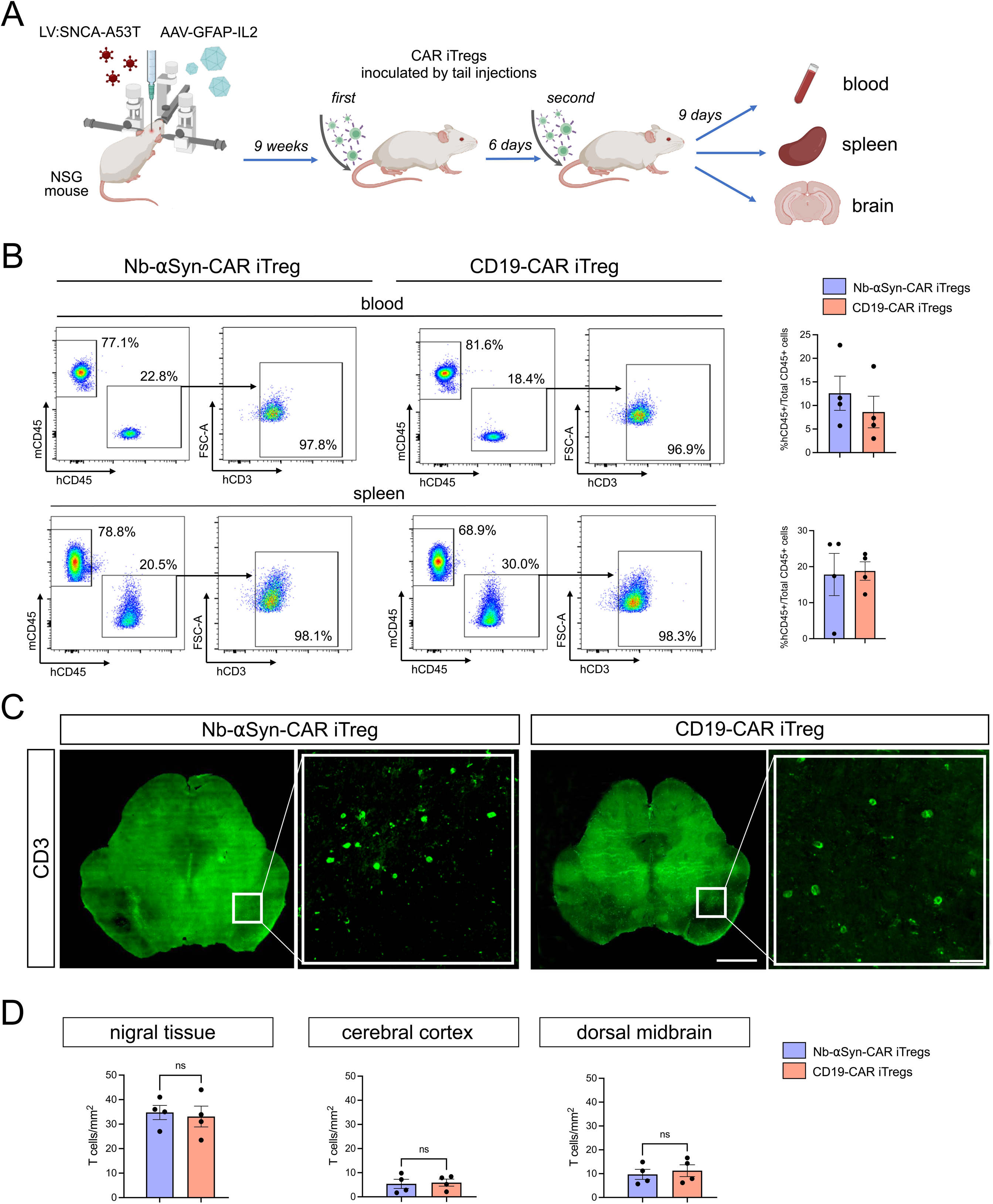
Biodistribution of CAR iTreg after systemic administration into NSG mice treated with LV-SNCA and AAV-GFAP-IL2. **A)** Schematic overview of the experimental design and its timeline to assess CAR iTreg homing after a double systemic infusion schedule. **B)** Gating strategy of FACS analysis used for the assessment of the percentage of human CD45^+^ and CD3^+^ cells in blood and spleen of animals infused with either Nb-aSyn-CAR iTregs or CD19-CAR iTregs and relative statistical analysis. 4 biological replicates per condition. **C)** Representative immunofluorescence images of coronal midbrain sections showing the entire midbrain and a higher-magnification view of the boxed region, highlighting CD3⁺ T-cell infiltration into the tissue of animals infused either with Nb-αSyn-CAR iTregs or CD19-CAR iTregs. Scale bars: 1 mm, 150 µm (inset). **D)** Quantification of the density of CD3+ T cells in nigral tissue, cerebral cortex and dorsal midbrain per unit area (1 mm^2^) in animals inoculated with either Nb-aSyn-CAR iTregs or CD19-CAR iTregs. n=4. All the data are presented as means ± SEM and were analyzed using *t*-student test. ns, *P* > 0.05.

### Nb-αSyn-CAR iTregs restrain neuroinflammation and αSyn pathology in PD mice

Next, Nb-αSyn-CAR iTregs or CD19-CAR iTregs were administered twice to mice (5×10⁶ cells/infusion/mouse) 8 weeks after LV:SNCA and IL2 viral transduction. Animals were euthanized 9 days later, and tissues were collected for the assessment of systemic biodistribution and brain neuroinflammation. (Figure 6A). CD19-CAR iTregs served as antigen-independent control cells to determine the specific effects mediated by the αSyn-targeting CAR. Flow cytometry analysis of hCD45 and hCD3 confirmed the presence of CAR iTregs in both the blood and spleen, with no significant differences in cell abundance between Nb-αSyn-CAR iTregs and CD19-CAR iTregs (Figure 6B). Brain sections were acquired by tile-scan imaging and reconstructed to quantify hCD3+ cells within the SNpc, cerebral cortex, and dorsal midbrain (Figure 6C). Notably, the highest number of infiltrating CAR iTregs was detected in the nigral tissue, indicating preferential homing to the brain region most severely affected by αSyn pathology (Figure 6D). However, no significant differences in CAR iTreg accumulation were observed between Nb-αSyn-CAR and CD19-CAR iTregs, suggesting that their recruitment to the substantia nigra is not driven by antigen recognition but is instead likely promoted by the highly inflammatory microenvironment induced by αSyn overexpression. Next, neuroinflammatory markers and αSyn pathology were assessed by quantitative immunofluorescence analysis. Prolonged αSyn overexpression markedly increased IBA1 and CD68 immunoreactivity, reflecting robust microglial activation (IBA1+ area: 3.9% ± 0.8%, 19.2% ± 1.9; CD68+ area: 0.5% ± 0.1, 6.3% ± 0.5, for AAV:IL-2 and AAV:IL2+LV:SNCA, respectively) (Figure 7A). This response was significantly attenuated in mice treated with Nb-αSyn-CAR iTregs, but not with CD19-CAR iTregs (IBA1+ area: 8.5% ± 1.3%, 20.2% ± 1.9%; CD68+ area: 1.7% ± 0.6%, 4.4% ± 0.7% for Nb-αSyn-CAR iTregs and CD19-CAR iTregs, respectively) (Figure 7A,C). Likewise, the increase in GFAP immunoreactivity associated with reactive astrogliosis showed a trend toward reduction only in the presence of Nb-αSyn-CAR iTregs, although this effect did not reach statistical significance (AAV:IL2: 6.9% ± 0.7%; AAV:IL2+LV:SNCA: 16.0% ± 1.4%; Nb-αSyn-CAR iTregs: 11.3% ± 1.0%; CD19-CAR iTreg: 16.9% ± 1.3%) (Figure 7B,C). Importantly, infiltration of Nb-αSyn-CAR iTregs also resulted in a significant reduction of both total human αSyn and phosphorylated αSyn (pS129αSyn), demonstrating a decrease in αSyn pathological burden (monomeric αSyn^+^ area: 1.0% ± 0.2% for AAV:IL2; 27.9% ± 6.5% for AAV:IL2+LV:SNCA; 9.4% ± 2.1% for Nb-αSyn-CAR iTregs; pS129αSyn^+^ area: 0% ± 0% for AAV:IL2; 3.2% ± 0.4% for AAV:IL2+LV:SNCA; 1.4% ± 0.2% for Nb-αSyn-CAR iTregs) (Figures 7A-C). In contrast, CD19-CAR iTregs failed to induce a comparable reduction in αSyn pathology (monomeric αSyn+ area: 19.9% ± 2.2%; pS129αSyn+ area: 2.8% ± 0.4%) (Figure 7A,C). To further characterize the anti-inflammatory effects of CAR iTreg treatment, an independent cohort of mice was analyzed at the same experimental endpoint, 9 days after the second and final CAR iTreg infusion. The substantia nigra was microdissected, and pro-inflammatory cytokines were quantified using a bead-based multiplex immunoassay. Notably, several key mediators of glial inflammation, including IL-1β, TNFα, and IL-6, were significantly reduced exclusively in mice receiving Nb-αSyn-CAR iTregs, whereas no significant changes were observed following treatment with CD19-CAR iTregs (Figure 7D). Collectively, these findings demonstrate that antigen-specific accumulation of Nb-αSyn-CAR iTregs within the inflamed brain elicits a potent antigen-specific anti-inflammatory response that mitigates glial activation and reduces αSyn pathology.

**Figure 7.**
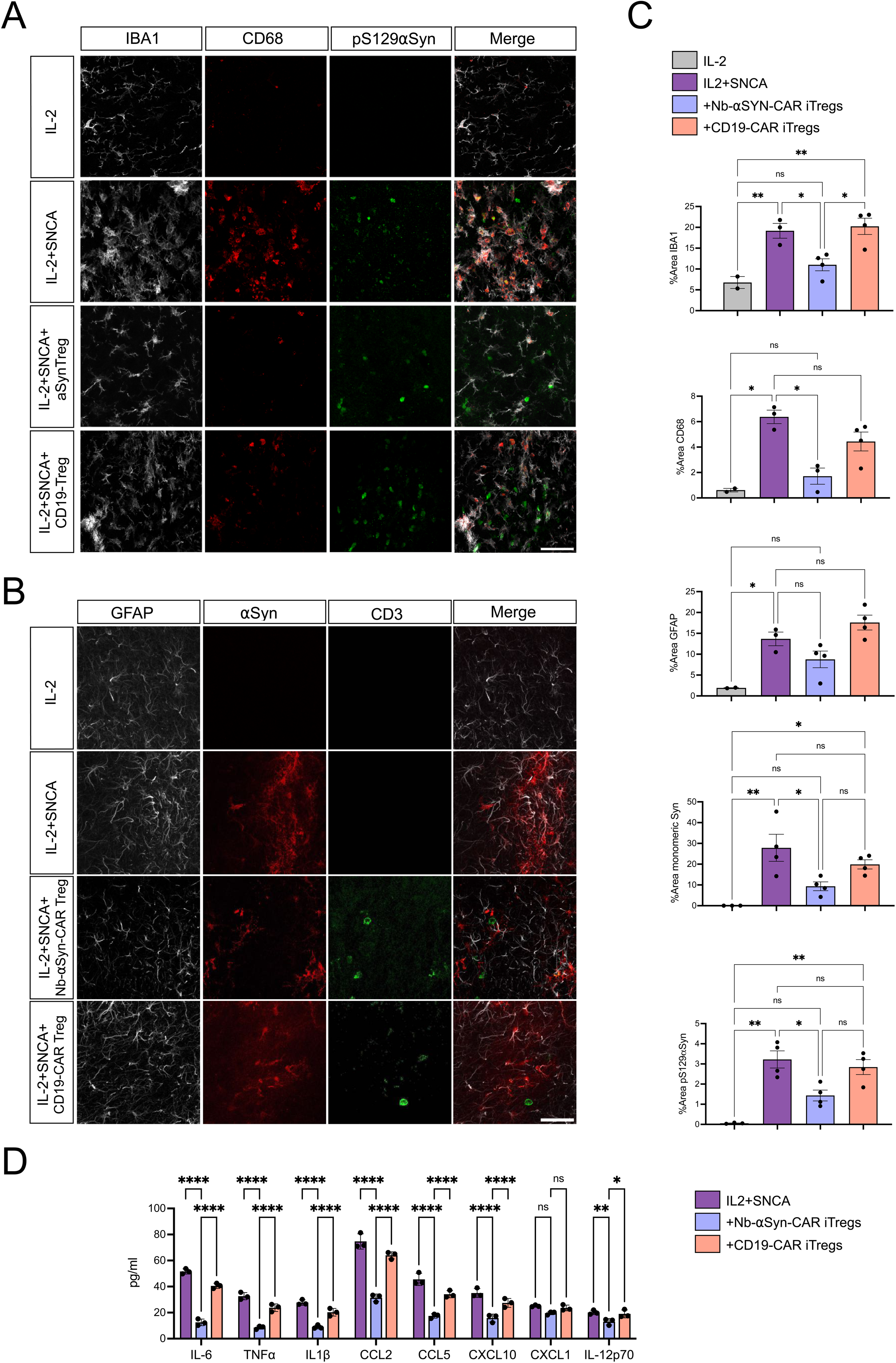
Nb-αSyn-CAR iTregs reduce neuroinflammation and αSyn accumulation burden in PD mice. **A)** Representative fluorescence photomicrographs showing IBA1 (gray), αSyn (red), pS129αSyn (green) and the merged fluorescence channels in comparable nigral areas across the four experimental conditions. Scale bar: 150 µm. **B)** Representative immunofluorescence images showing GFAP (gray), αSyn (red), CD3 (green) and the merged fluorescence channels in comparable regions of *substantia nigra* across the four experimental conditions. Scale bar: 150 µm. **C)** Bar graphs showing the quantification of the immunofluorescence-positive area for IBA1, CD68, GFAP, αSyn, and pS129αSyn. Data were obtained from four independent fields of view per animal (3 biological replicates). All data are presented as the means ± SEM and were subjected to analysis using one-way ANOVA followed by Tukey’s multiple comparisons test. \**P* < 0.05; \*\*\**P* < 0.001; \*\*\*\**P* < 0.0001; ns, *P* > 0.05. **D)** Bar graph with quantification of the cytokines in the dissected nigral tissue in the different experimental conditions as obtained by the bead-based multiplex immunoassay. Data are presented as means ± SEM and were analyzed using one-way ANOVA followed by Tukey’s multiple comparisons test. \**P* < 0.05; \*\**P* < 0.01; \*\*\*\**P* < 0.0001, ns, *P* > 0.05.

## Discussion

In this study, we engineered human αSyn-CAR iTregs capable of selectively sensing αSyn preformed fibrils (PFFs) and activating their suppressive program in an antigen-dependent manner. In the presence of human IL-2, αSyn-CAR iTregs persisted for several days within the degenerating nigral tissue and exerted a strong anti-inflammatory effect, dampening both microglial and astrocytic reactivity. Notably, *in vitro* selective antigen-dependent activation was achieved only with the NbSyn87-based CAR, whereas the scFv-based construct displayed non-antigen-specific activation in the absence of cognate antigen. This finding is consistent with previous studies showing that scFvs are prone to misfolding and self-association on the T-cell surface, leading to ligand-independent tonic signaling and progressive T-cell exhaustion (Long et al., 2015). By contrast, nanobodies possess intrinsic structural features that make them particularly attractive for CAR engineering. Their compact single-domain architecture enhances stability and solubility while reducing the propensity for receptor aggregation caused by inter-domain interactions (Boisgard et al., 2026). Moreover, their small size facilitates access to cryptic epitopes that are often inaccessible to conventional antibodies (Ditlev et al., 2014), including conformational epitopes embedded within the densely packed architecture as that featured by αSyn aggregates. We demonstrated that the Nb-based CAR was selectively activated by aggregated αSyn conformers, while exhibiting minimal activation in response to monomeric αSyn. This result is particularly noteworthy considering that we showed that the parental NbSyn87 nanobody recognizes both monomeric and aggregated αSyn species on brain sections (Figure 2), consistent with its previously reported *in vitro* binding properties (Guilliams et al., 2013). These data therefore suggest that incorporation of NbSyn87 into the CAR architecture confers a functional selectivity for aggregated αSyn that is not obvious from the binding profile of the nanobody alone. A likely explanation is the multivalent presentation of epitopes on αSyn PFFs, which promotes CAR crosslinking and clustering at the T-cell surface. Such multivalent engagement is a well-established requirement for efficient CAR signaling, as robust downstream activation typically depends on the simultaneous engagement of multiple CAR molecules by repetitive antigenic epitopes (Wu et al., 2020). In contrast, monomeric αSyn lacks the multivalency necessary to induce sufficient receptor clustering and therefore fails to trigger meaningful CAR activation (Wu et al., 2020). This selective responsiveness is especially important given the physiological diversity and compartmentalization of αSyn. Even though monomeric αSyn is primarily intracellular and not readily accessible for CAR interaction, the ability to specifically target pathological extracellular aggregates optimizes therapeutic precision. By targeting aggregated αSyn fibrils that accumulate within diseased brain regions, CAR Tregs can preferentially exert their immunomodulatory activity where neuroinflammation and neurodegeneration are most prominent, while sparing physiological monomeric αSyn. Such selectivity is essential, as monomeric αSyn is expressed by several peripheral cell types, including neuroendocrine cells (e.g., pancreatic β-cells and enteroendocrine cells) and hematopoietic cells, with mature erythrocytes representing the largest peripheral reservoir of αSyn (Barbour et al., 2008). Consequently, restricting CAR activation to aggregated αSyn species is a key safety feature that may limit off-target activation and mitigate undesired systemic immunosuppression. To evaluate the therapeutic relevance of human Nb-αSyn-CAR iTregs, we first developed and validated an immunodeficient mouse model of synucleopathy based on lentiviral-mediated SNCA-A53T overexpression, which recapitulated pathological αSyn accumulation and pronounced glial reactivity while remaining permissive to human cell xenotransplantation. Despite this favorable environment, human iTregs displayed impaired survival following transplantation. Notably, astrocyte-directed viral expression of human IL-2 robustly rescued iTreg persistence, highlighting the critical role of species-matched IL-2 signaling in supporting engraftment within the murine CNS. This finding is consistent with evidence that murine IL-2 only inefficiently sustains human Treg biology (Robert et al., 2025). In addition, NSG mice are devoid of lymphocytes, the major physiological source of IL-2, and the healthy brain expresses only low levels of this cytokine (Yshii et al., 2022). Together, these features render the murine CNS a harsh environment for the survival and maintenance of xenotransplanted human Tregs, requiring human IL-2 AAV-driven expression.

Our findings provide strong evidence that iTregs recruited to pathological brain regions can mount a robust local immunomodulatory response. iTreg administration markedly reduced the inflammatory state of both microglia and astrocytes, leading to a prominent decrease in pro-inflammatory cytokines, activation markers, and reactive morphological features. Given the central role of neuroinflammation in amplifying αSyn pathology and enhancing DAergic neuronal vulnerability, these results suggest that Treg-based immunotherapy may interrupt a key pathogenic feed-forward mechanism underlying Parkinson’s disease progression (Duffy et al., 2018; Bido et al., 2024). Corroborating this hypothesis, both monomeric and aggregated αSyn species were reduced in the nigral tissue infiltrated by Nb-αSyn-CAR iTregs, suggesting enhanced clearance of αSyn by resident glial cells. This effect may result from the attenuation of glial inflammatory activation, which is expected to improve the phagocytic and degradative functions of these cells toward pathological αSyn species. Whether such immune modulation ultimately translates into delayed neurodegeneration and sustained preservation of nigrostriatal circuitry remains an important question for future investigation.

Strengthening Treg activity has a strong rational in PD where their suppressive functions appear to be affected in at least some cohorts of patients (Kustrimovic et al., 2018; Álvarez-Luquín et al. 2019) and Th1 pro-inflammatory cells are more generally increased (Wang et al., 2021; Capelle et al., 2023; Moquin-Beaudry et al., 2025). In addition, it has been shown that depletion of Tregs in PD mice accelerated neurodegeneration, confirming their key role in maintaining CNS immune homeostasis (Li et al., 2021). Importantly, adoptive transfer of CD4^+^CD25^+^ Tregs mitigates DAergic neuronal loss and decreases microglial activation and lymphocyte infiltration in MPTP and αSyn treated mice, indicating that Tregs can reduce both innate immune effector functions and downstream neuronal injury (Reynolds et al., 2007; 2010; Huang et al., 2020; Li et al., 2021; Badr et al., 2022). However, the clinical translation of adoptive Treg cell transfer or systemic expansion strategies is significantly hindered by several pharmacological and biological hurdles. In fact, clinical trials conducted with expanded polyclonal Tregs proved to be safe, but resulted in unsatisfactory clinical improvements, possibly due to the low amount of antigen-specific regulatory cells in the infused product and the lack of tissue-targeted specificity (Raffin et al., 2020; Bender et al., 2024). CAR engineering might overcome these limitations by enhancing Treg immunosuppressive functions exclusively in the presence of the antigen. Along this line, some of us recently showed that Fox19-CAR iTregs efficiently suppressed proliferation and activity of B cells *in vitro*, restricting autoantibody generation in a humanized mouse model of the autoimmune disease systemic lupus erythematosus (Doglio et al., 2024). In the present work we further extended the wide applicability of a CAR iTreg cellular product for the treatment of a neuroinflammatory neurodegenerative disease.

In summary, this study establishes a robust platform for generating functional αSyn-specific CAR iTregs with precise antigen selectivity and prominent suppressive capacity, and highlights essential considerations for their translational application in PD. The integration of human immune cells, optimized genetic engineering, and physiologically relevant models offers a clear path toward clinical development of next-generation antigen-specific immunotherapies for PD.

## Materials and Methods

### CAR construct design

The CAR constructs utilized are based on a second-generation architecture incorporating CD28 transmembrane and costimulatory domain, and intracellular CD3ζ stimulatory domain. Furthermore, all constructs incorporate a long spacer derived from the wild-type nerve growth factor receptor (NGFR), previously engineered to enable surface detection (Casucci et al., 2018). The LV Foxp3_CD19 CAR construct consists in a unidirectional LV containing the human FoxP3 gene and the anti-CD19 CAR linked by a thosea-asigna virus 2A (T2A) peptide and controlled by a hPGK promoter, as previously described by Doglio and colleagues (Doglio et al., 2024). The LV Foxp3_αSyn-NB-CAR was generated by substituting the anti-CD19 scFv with an antigen-binding domain derived from the NbSyn87 nanobody (NB), which recognizes both aggregated and monomeric forms of αSyn (Guilliams et al., 2013). While the αSyn-scFv-CAR embeds a scFv derived from the VH and VL domains of the anti-Syn antibody 306C7B3 (Düchs et al., 2023), connected via the Whitlow linker (Whitlow et al., 1993).

### Lentiviral vector production

VSVg-pseudotyped, replication-incompetent lentiviral particles were produced in HEK293T cells by calcium phosphate transfection as previously described (Bido et al., 2024). Briefly, 7.5×10⁶ HEK293T cells were plated on 150 mm dishes 24h prior to transfection. Supernatant was collected at 30h, filtered through a 0.22 µm cellulose acetate membrane, and ultracentrifuged at 50,000×g for 2 h at room temperature. Pellets from two dishes were resuspended in 80 µL PBS and stored at −80°C. Viral titer was determined by qPCR using the LV900 kit (Gentaur).

### Isolation and processing of CD4⁺ T cells

Primary human CD4⁺ T cells were isolated from healthy donor buffy coats (written informed consent, San Raffaele Institutional Ethics Committee) by negative selection using RosetteSep Human CD4⁺ T Cell Enrichment Cocktail (#15022, STEMCELL Technologies) over Histopaque-1077 density gradient medium (Sigma-Aldrich, H8889). Enriched cells were washed twice with PBS/10% FBS, counted, and either used immediately or stored at −80°C.

Cells were cultured in X-VIVO 15 (Lonza, BW04-418Q) supplemented with 10% FBS and 100 µg/mL Primocin (InvivoGen). Activation was performed with αCD3/CD28 coated Dynabeads (Gibco, 11131D) and 200 U/mL human IL-2 (Biolegend, 791906) for 48 h prior to LV transduction using MOI 100. Three days post-transduction, medium was supplemented with 500 U/mL hIL-2 and 500 nM rapamycin (Merck, R8781) to support Treg expansion. Cells were restimulated with αCD3/CD28 beads on day 9 and rested from day 14; cryopreservation was performed on day 21.

### Flow cytometry

CAR iTreg transduction efficiency and phenotype were assessed by flow cytometry. Cells were stained for 10min at room temperature with a Live/Dead viability dyes (Invitrogen, L34975; Invitrogen, L3224) and for 10 min at 4°C with surface antibodies against CD4 (Miltenyi Biotec, 130-126-322 or 130-113-226), CD25 (Miltenyi Biotec, 130-113-849 or Invitrogen, 25-0257-42), LNGFR (Miltenyi Biotec, 130-112-790 or BD Biosciences, 564580), HLA-DR (Biolegend, 307618), CTLA-4 (Biolegend, 369610), GARP (BD Biosciences, 583958). Following fixation/permeabilization with the FOXP3 Transcription Factor Staining Buffer Set (Invitrogen, 00-5523-00), intracellular staining was performed for 30min at room temperature with anti-FOXP3 (Invitrogen, 17-5773-82) and anti-HELIOS (Biolegend, 137216) antibodies.

For flow cytometric analysis of blood and spleen, on the day of sacrifice mice were deeply anesthetized and blood was collected by retroorbital-bleed into EDTA-coated tubes, followed by transcardial perfusion with saline. Spleens were then harvested and mechanically dissociated through 70 µm cell strainers to obtain single-cell suspensions. Red blood cells in both blood and splenocyte suspensions were lysed using ACK (ammonium-chloride-potassium) lysing buffer for 5 min at room temperature, followed by washing with FACS buffer (PBS, 0.5% BSA, 2mM EDTA). Cells were stained for 10 min at 4°C with antibodies against surface markers: mCD45 (Invitrogen, 45-0451-82), hCD45 (Miltenyi Biotec, 130-110-637), hCD3 (Miltenyi Biotec, 130-113-698). After staining, cells were washed twice with FACS buffer and fixed with 1% PFA (Merck) prior to acquisition. Samples were acquired on a BD FACSCanto II flow cytometer (BD Biosciences), and data were analyzed using FlowJo software.

For flow cytometric analysis of microglia-like cells, they were detached with accutase, washed with PBS, stained with a Live/Dead viability dye as previously described, and then with antibodies against human CD45 (Miltenyi Biotec, 130-110-637), CD14 (Miltenyi Biotec, 130-110-521), TMEM119 (Biolegend, 853315), CD86 (Miltenyi Biotec, 130-116-267), HLA-DR (Biolegend, 307618), CD206 (Biolegend, 321110), CD163 (BD Biosciences, 563697).

### RNA extraction and real time qPCR

Total RNA was extracted using NucleoZol (Carlo Erba) following manufacturer’s instructions. 1 µg of RNA was reverse transcribed using ImProm-II™ Reverse Transcription System (Promega). For quantitative real time PCR (RT-qPCR), Titan HotTaq EvaGreen qPCR mix (BioAtlas) was used and expression levels were normalized respect to 18S expression. The results were reported as the 2^−ΔΔCt^. Primers for gene expression analysis are listed in Supplementary Table 1.

### Lentiviral vector copy number

Lentiviral vector copy number (VCN) was determined by droplet digital PCR (ddPCR) to quantify integration of the lentiviral constructs in transduced induced regulatory T cells (iTregs). Primers and a FAM-labeled hydrolysis probe were designed against a sequence common to all three lentiviral constructs used to generate αSyn-NB, αSyn-scFv, and CD19-CAR iTregs, enabling a single assay applicable across constructs, while THNSL2 served as the HEX-labeled single-copy reference gene for genomic DNA normalization. Reactions contained ddPCR Supermix for Probes No dUTP (Bio-Rad, #1863024), and droplets were generated using the QX200 Droplet Generator (Bio-Rad, #1864002). Following amplification by standard thermal cycling, reading was performed on the QX200 Droplet Reader (Bio-Rad, #1864003). Vector copy number per diploid genome was calculated using QuantaSoft software (Bio-Rad).

### Antigen-binding domain validation

The anti-αSyn NB and scFv were validated for antigen recognition on brain slices overexpressing α-Syn. Each domain was expressed as a fusion protein with a human IgG Fc region, produced in HEK293T cells transduced with the corresponding LV and selected with puromycin. Fc-fusion proteins were secreted into PRO293a serum-free medium (Lonza), collected at 72 h, and filtered. Supernatants were used as primary antibody substitutes during immunofluorescence (3% BSA, 0.3% Tween-20).

### In vitro assays

For proliferation assays, cells were labeled with eFluor 450 or eFluor 670 Cell Proliferation Dyes (Thermo Fisher Scientific, 65-0842-85 and 65-0840-85) at 1:1000 in PBS following vendor instructions.

For activation assays, 5×10⁴ iTreg cells were seeded in 96-well round-bottom plates. Cells were cultured for five days in the following conditions: unstimulated (no IL-2); IL-2 alone (500 U/mL); and IL-2 with αCD3/CD28 beads (1:1 bead-to-cell ratio), αSyn monomers (1 µM), or αSyn PFF (1 nM). αSyn monomers and PFF were kindly provided by Drs. A. De Simone’s and Professor R. Melki’s laboratories, respectively.

For polyclonal suppression assays, Tconv (5×10⁴/well) were co-cultured for five days with serially diluted Tregs at Treg:Tconv ratios of 2:1, 1:1, 1:2, 1:4, and 1:8 in αCD3/CD28 bead-stimulated conditions (1:1). No exogenous IL-2 was added. Control conditions included Tconv and Tregs cultured separately, with or without stimulation.

For antigen-specific suppression assay, cells were separately simulated prior to assay co-culture. αSyn-NB- and CD19-CAR iTregs were pre-stimulated for 48 h with 500 U/mL IL-2 and 1 nM PFF. Negative controls received no stimulus; positive controls received αCD3/CD28 beads (1:1). Tconv were pre-stimulated for 24h with αCD3/CD28 beads (1:10) and 200 U/mL IL-2. After pre-stimulation, beads were magnetically removed, and cells were washed and counted before co-culture setup. Tregs were seeded in serial dilutions, Tconv added at 5×10⁴/well, and PFF included in wells corresponding to αSyn-stimulated Treg conditions. After 48h of co-culture, supernatant was collected for ELISA, cells were washed, stained with a viability dye, fixed, and acquired on a CytoFlex S cytometer (Beckman Coulter). Percent suppression was calculated as following: Percent Suppression = [(% proliferation of Tconv alone - % proliferation of Tconv with Tregs) / % proliferation of Tconv alone] × 100.

### Induced Microglia-like cells (iMGCs)

iMGCs were differentiated from monocytes obtained from healthy donor buffy coats. Total buffy coats were diluted 1:1 with PBS and layered onto Histopaque-1077 density gradient medium (Sigma-Aldrich, H8889) and centrifuged to obtain peripheral blood mononuclear cells (PBMCs). Monocytes were further purified by density-gradient centrifugation over 46% osmolarity-adjusted Percoll. The monocyte-enriched interphase was collected and plated in serum-free RPMI to allow selective adhesion for 20 min. To generate iMGCs, adherent monocytes were subsequently cultured in complete RPMI supplemented with 10% FBS, 1% penicillin-streptomycin, 1% L-glutamine, 10 ng/mL human GM-CSF (Miltenyi Biotec, 130-093-865) and 100 ng/mL human IL-34 (Miltenyi Biotec, 130-108-977), with medium replaced every 4 days for up to 14 days of differentiation.

For iMGC–Treg co-cultures, CAR iTreg cells were primed for 48 h with either PFFs or αCD3/CD28 beads. CAR iTreg and iMGCs were then co-cultured at a 1:1 (Treg:iMGC) ratio in the presence of PFFs for 24h. Then, inflammatory stimuli consisting of 20ng/ml IFNγ and 50ng/ml LPS were added for 6 h for RNA extraction and for 24 h for flow cytometry analyses. At the end of the assay, CAR iTreg cells were removed as the non-adherent fraction, while iMGCs activation was evaluated by RT-qPCR analysis and flow cytometry.

### ELISA

Culture supernatants from activation assays were analyzed for IL-10 production using a human IL-10 ELISA kit (Invitrogen, EHIL10) following the manufacturer’s instructions. Briefly, samples and standards were incubated 2 h at room temperature in pre-coated plates, followed by HRP-conjugated detection antibody (1 h), TMB substrate (10 min, dark), and stop solution. Absorbance was measured at 450 nm with 550 nm reference subtraction; IL-10 concentrations were interpolated from a standard curve by linear regression.

### Animals

NOD.Cg-*Prkdc*^scid^ *Il2rg^tm1Wjl^*/SzJ (NSG) mice were obtained from Jackson Laboratories and housed at the San Raffaele Hospital Animal Facility under a 12h light/dark cycle, controlled temperature (25°C) and humidity (50–60%), with ad libitum access to food and water. All procedures were conducted in accordance with protocols approved by the Institutional Animal Care and Use Committee and reported to the Italian Ministry of Health in compliance with EU Directive 2010/63/EU.

For stereotaxic injections, mice were anesthetized with isoflurane. LVs (1×10⁹ TU/mL; 34-gauge needle) or T cells (5×10⁴ cells; 26-gauge needle, adjusted dorso-ventral coordinate) were delivered into the *substantia nigra pars compacta* at a volume of 1.5µL and flow rate of 0.25µL/min using the following injection coordinates (antero-posterior / medio-lateral / dorso-ventral, in mm from bregma): −3.9 / ±1.9 / −4.9 for LV injections and −3.9 / ±1.9 / −4.7 for T cell injections. Control animals received an equivalent volume of empty viral capsid preparation or PBS, respectively.

PD pathology was induced by bilateral intracranial injection of LV SNCA^A53T^. Engineered cells were infused intravenously (5×10⁶ cells) at 8 weeks post-LV injection, with terminal collection of peripheral blood, spleen and brain tissue at 9 days post-final infusion.

### Immunofluorescence

For cellular immunofluorescence, CD4⁺ T cells (1.5×10⁵) were cytocentrifuged onto glass slides (Cytospin 4, Thermo Fisher Scientific) and fixed with 4% PFA. For tissue immunofluorescence, mice were transcardially perfused with saline followed by 4% PFA, brains were post-fixed overnight in 4% PFA, cryoprotected in PBS/20% sucrose (24 h), flash-frozen, and sectioned at 50 µm on a cryostat; free-floating sections were additionally treated with 3% H₂O₂/10% methanol (10 min) and permeabilized with 2% Triton X-100/PBS (20 min). Both cell and tissue preparations were then blocked (1 h, room temperature; PBS, 3% BSA, 0.3% Tween-20) and incubated overnight at 4°C with primary antibodies directed against CD3 (Bio-Rad, MCA1477), NGFR (Cell Signaling, 8238T), αSyn (Thermo Fisher, 180215), pS129αSyn (Abcam, ab51253), CD68 (Abcam, ab53444), GFAP (Abcam, ab4674), Iba1 (DBA, 234009), or TH (Millipore, MAB318). Secondary antibodies and Hoechst were incubated for 1 h at room temperature, and coverslips were mounted with Dako fluorescence mounting medium (S3023).

### Acquisition and quantification of immunofluorescence images

Immunofluorescence images were acquired using a Leica TCS SP8 laser-scanning confocal microscope equipped with a 40× and 63× objective. Wide-field images were acquired using an ImageXpress High-Content Micro Confocal Imaging System. Image analysis was performed in ImageJ using automated macros developed for each experimental workflow. To ensure unbiased quantification, all images were analyzed in a blinded manner using a fixed fluorescence intensity threshold that was defined a priori and consistently applied to all images within each experimental set.

### Quantification of CD3-positive cells in the substantia nigra

Tissue sections (50um thickness) were imaged using an ImageXpress high-content imaging system equipped with a 20× Plan Apo Lambda dry objective (numerical aperture, 0.75). For each section, a region of interest (ROI) encompassing the *substantia nigra* was manually delineated. CD3-positive (CD3⁺) cells within the selected ROI were identified and counted for each experimental group. The area of each ROI was measured and converted to mm^2^. CD3⁺ T-cell density was then calculated by dividing the number of CD3⁺ cells by the corresponding ROI area and was expressed as cells/mm².

### Statistical Analysis

Flow cytometry data were analyzed with FlowJo v10.10.0 (BD Biosciences). Statistical analyses and graph generation were performed in GraphPad Prism v10.0.0. Data are presented as mean ± SEM unless otherwise stated. Two-group comparisons were performed with unpaired t-tests; multiple comparisons were analyzed by one-way or two-way ANOVA with appropriate post hoc tests. P < 0.05 was considered statistically significant.

## Lead contact

Further information and requests for resources and reagents should be directed to and will be fulfilled by the lead contact, Vania Broccoli.

## Materials availability

Resources and materials will be shared by the lead contact upon reasonable request.

## Acknowledgements

We are grateful to S. Gregori and all members of the Broccoli’s lab for helpful discussion. We acknowledge the FRACTAL and ALEMBIC core facilities for expert supervision in flow-cytometry and confocal imaging, respectively. Some illustrations used in this work were created in BioRender (https://BioRender.com). This work was supported by ERC (#101199846-CARgiver) to V.B.

## Author contributions

Conceptualization, A.C., S.M. and V.B.; methodology, A.C., C.B.B., M.D., G.S.S., R.M., M.C. and C.B.; investigation, A.C., M.N., G.R., G.S.S. and S. M.; data curation, A.C., M.N., S.M. and V.B.; writing – original draft preparation, A.C., S.M. and V.B.; writing – review & editing, A.C., M.N., M.D., M.C., C.B., S.M. and V.B.; visualization preparation, A.C., M.N., S.M. and V.B.; supervision, project administration, and funding acquisition, V.B.

## Supplementary Materials

**Supplementary Figure 1.**
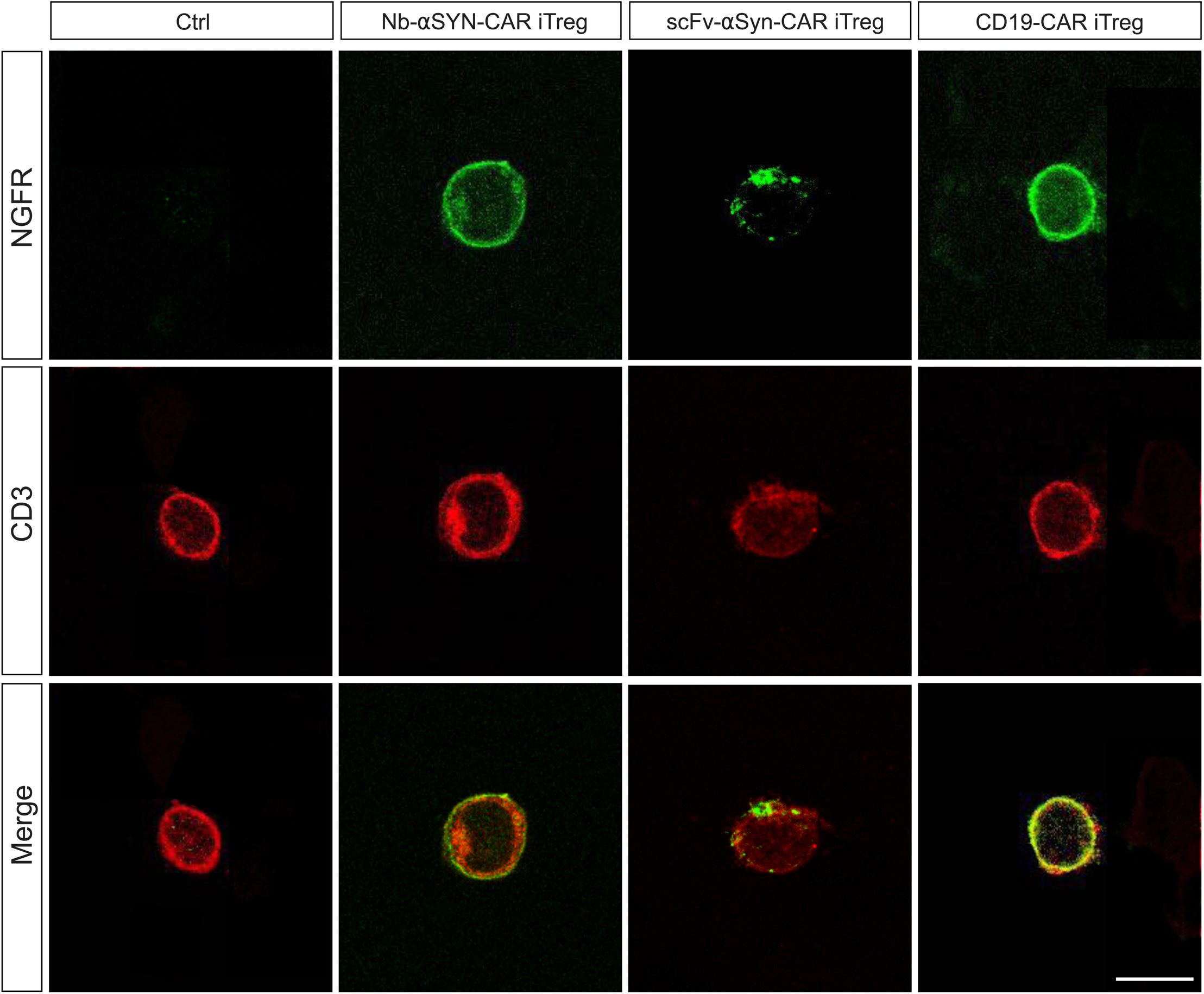
Membrane localization of CAR constructs in engineered CAR iTregs. Representative confocal microscopy images showing cell surface expression and co-localization of NGFR (green) and CD3 (red) in untransduced control cells (Ctrl), Nb-αSYN-CAR, scFv-αSYN-CAR, and CD19-CAR iTregs. Merged images (bottom row) demonstrate cell membrane co-localization of NGFR with CD3 (yellow overlay). Scale bar: 30 µm.

**Supplementary Figure 2.**
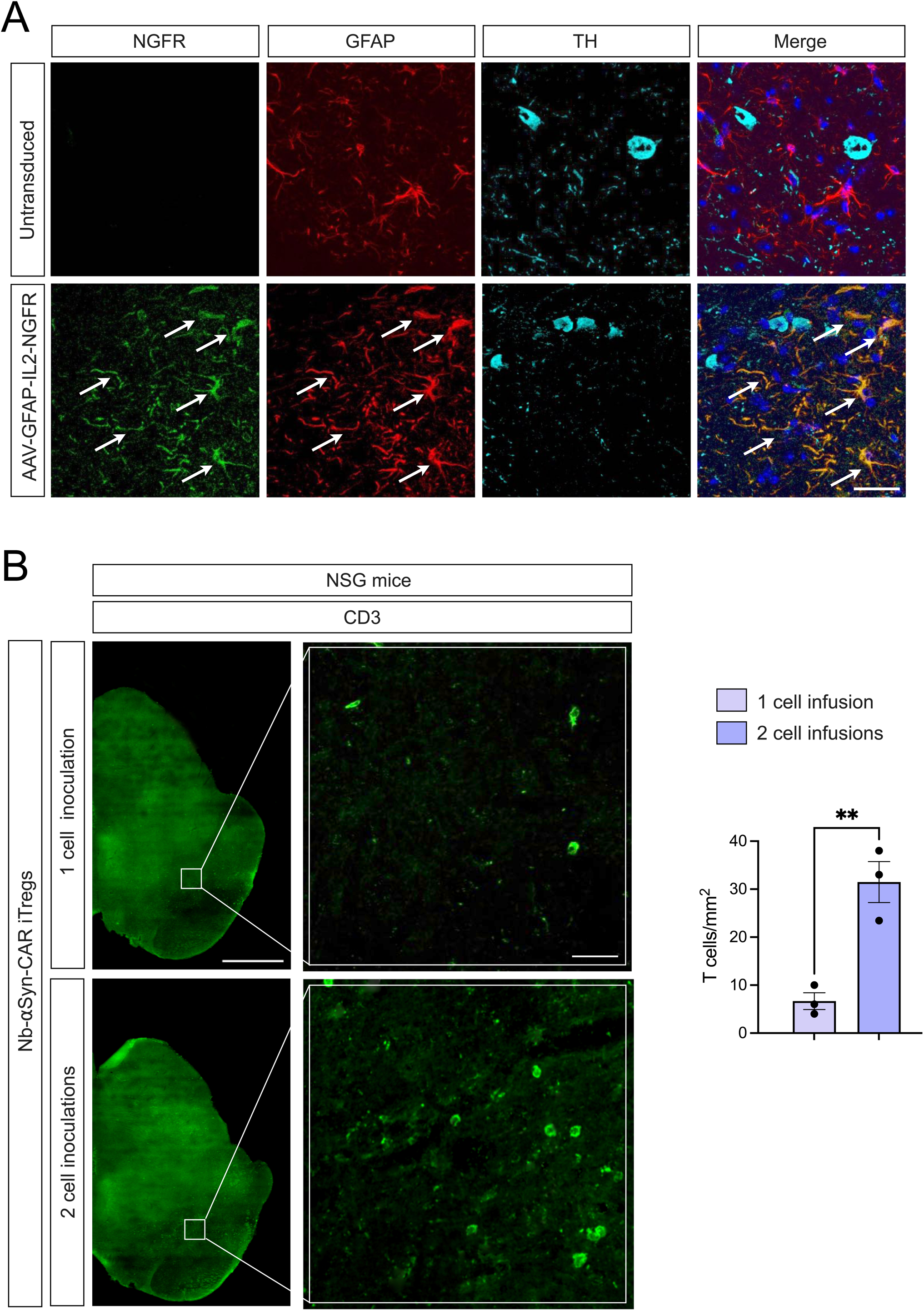
Trasduction pattern of AAV-GFAP-IL2-NGFR and T cell brain infiltration quantifications. **A)** Representative immunofluorescence images of comparable regions of the *substantia nigra* stained for NGFR (green), GFAP (red), and TH (cyan), together with the merged fluorescence channels. Arrowheads indicate the extensive co-localization of NGFR with GFAP, consistent with astrocyte-specific NGFR expression driven by the AAV-GFAP vector. Images are representative of three independent biological replicates. Scale bar: 150 μm. **B)** Representative low- and high-magnification immunofluorescence images of CD3+ T cells infiltrating the brain following one or two intravenous administrations via tail vein injection, together with the corresponding quantification. Scale bars: 1 mm, 150 μm (inset). 3 biological replicates. Data are presented as means ± SEM and were analyzed using *t*-student test. \*\**P* < 0.01.

**Supplementary Table 1.**
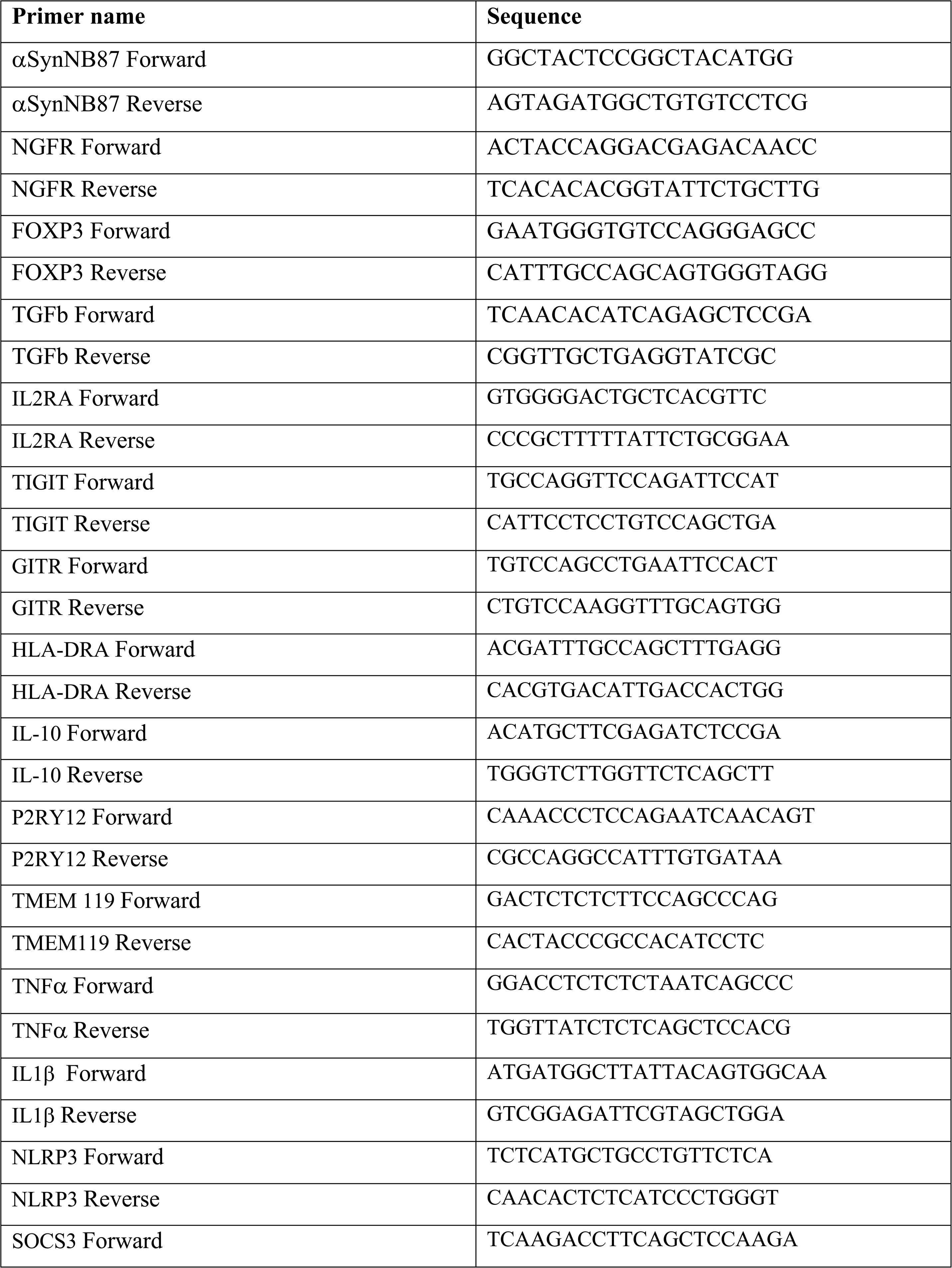
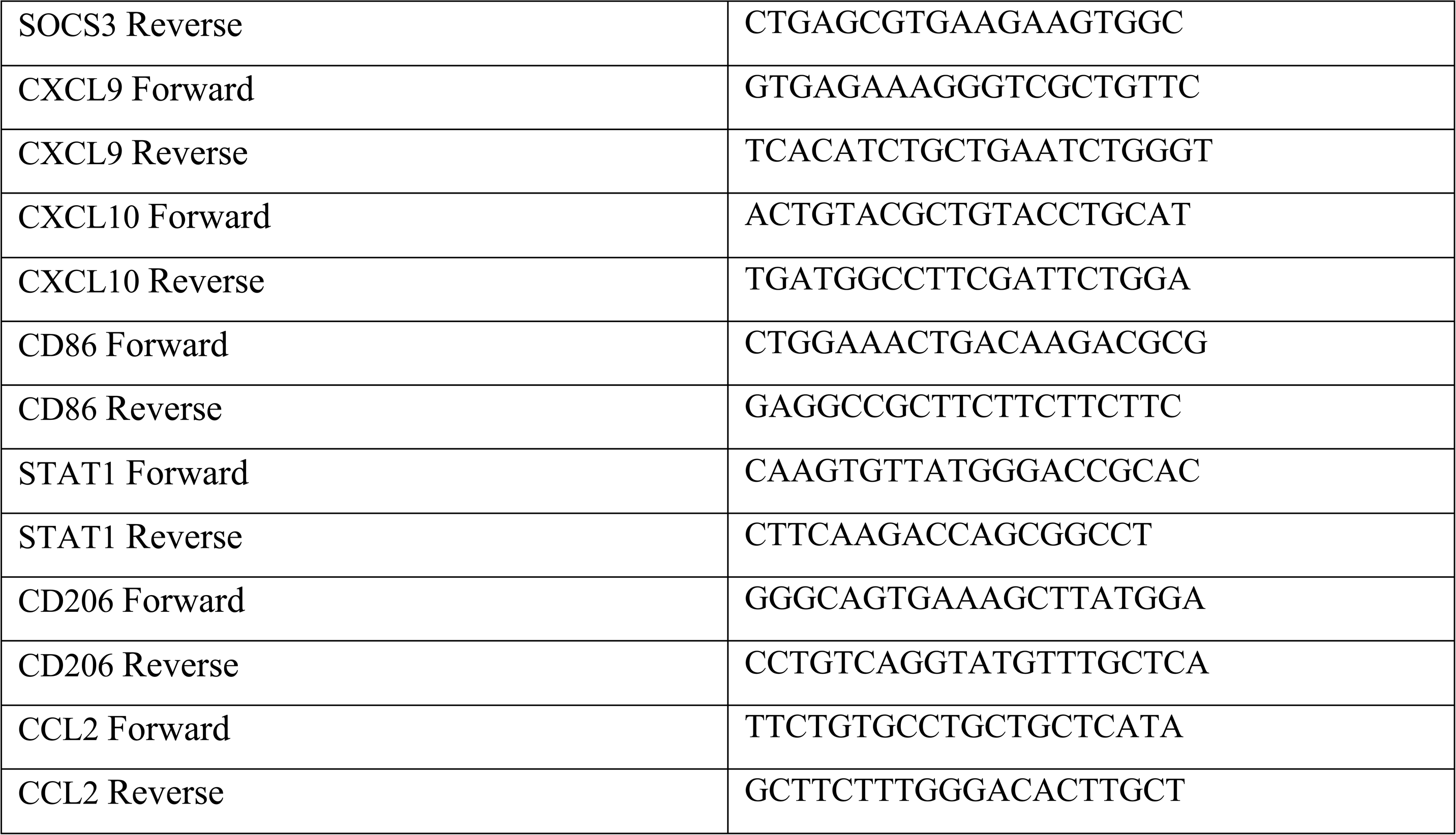
List of the primers used in this study for gene expression studies.

